# The phytolongin AtPhyl2.1 is involved in cell plate formation and root development

**DOI:** 10.64898/2025.12.15.694276

**Authors:** Valerie Wattelet-Boyer, Matthieu Buridan, Yuri L. Negroni, Chiara Mafficini, Franziska Dittrich-Domergue, Lilly Maneta-Peyret, Emily Breeze, Michela Zottini, Elide Formentin, Francesco Filippini, Lysiane Brocard, Patrick Moreau

## Abstract

SNAREs are critical elements of the membrane trafficking machinery with a wide variety of functionality across this family of proteins. Phytolongins are a recently identified subfamily of longins which possess the typical longin domain but lack a SNARE motif. Phytolongins have an ubiquitous tissue expression in Arabidopsis and are distributed throughout the secretory pathway. We focused on Phytolongin 2.1 (PHYL2.1) which localizes to the endoplasmic reticulum, and observed a strong root growth phenotype in the loss-of-function *Atphyl2*.*1-1* mutant. We demonstrate that whilst cell elongation efficiency was not affected in the mutant, cell division was significantly reduced. The resulting decrease in root length in the *Atphyl2*.*1-1* mutant is explained by a smaller number of cells which then elongate to enable root growth. Root apical meristem architecture of *Atphyl2*.*1-1* and another mutant *Atphyl2*.*1-2* was disturbed and distances from the root quiescent center to the transition zone and the first areas of mis-organized cells were affected in both mutants. Investigation of the SNARE AtKNOLLE revealed significant perturbation of *Atphyl2*.*1-1* cell plate formation in the mis-organised areas. Our results provide a first characterization of the phytolongin AtPHYL2.1 which appears involved in root cell plate formation, root cell division and therefore root development.

**Highlight:** The phytolongin AtPHYL2.1 significantly affects the efficiency of cell plate formation and root development in *Arabidopsis thaliana*.

## Introduction

SNAREs (Soluble N-ethylmaleimide-sensitive factor Attachment REceptors) are small membrane proteins which are widely involved in membrane trafficking processes and this is naturally the case in root growth and development (Luo et al., 2022). There is a huge diversity of SNAREs which is explained by their requirements in many physiological functions during plant life (Martinière and Moreau, 2020). In this diversification, the SNARE Longins (characterized by a N-terminal longin domain and the SNARE motif [Rossi et al., 2004]) represent a large superfamily (De Franceschi et al. 2014) which are essential and present in all eukaryotes.

A subfamily of four non-SNARE longins (containing a N-terminal longin domain but lacking a typical SNARE motif) has been identified in land plants (Vedovato et al., 2009). Structural modeling of the longin domain indicated that it was orthologous to the longin domain of the SNARE VAMP7 (Vedovato et al., 2009). In *Arabidopsis thaliana*, the four identified non-SNARE longins were named phytolongins and abbreviated as PHYL1.1, PHYL1.2, PHYL2.1 and PHYL2.2 (De Marcos Lousa et al, 2016).

Phytolongins have a rather ubiquitous tissue expression with PHYL2.1 and PHYL2.2 being exclusively localized to the endoplasmic reticulum (ER) network; PHYL1.2 being associated with the Golgi membranes and PHYL1.1 being mainly localized at the plasma membrane but also distributed to some extent in the Golgi, the trans-Golgi network and prevacuolar compartments (De Marcos Lousa et al, 2016). It was determined that a surface exposed YF motif (present in the □1-□3 region of the longin domain) was critical for the ER export of PHYL1.1 (Y48F49 motif) and that such a motif (Y50F51) was potentially also required for the ER export of PHYL1.2 (De Marcos Lousa et al, 2016).

When considering the role played by the SNARE domain in subcellular trafficking, its absence in phytolongins might suggest that these longins may not be engaged in any trafficking process. However, the association between the longin domain and subcellular trafficking is a common rule (De Franceschi et al., 2014), and even some longins which are missing a SNARE domain (mammalian Sec22b homologues Sec22a/c) are still involved in subcellular trafficking (Rossi et al., 2004).

In this study, we investigated the potential role of the ER-localised AtPHYL2.1in root cell growth. We identified a strong root growth phenotype in the *Atphyl2*.*1-1* loss-of-function mutant (RATM53-3274-1) and showed that cell elongation efficiency was in fact not affected in the *Atphyl2*.*1-1* mutant but that cell division was clearly disturbed. The decrease in root length in the *Atphyl2*.*1-1* mutant could be at least partly explained by a lower number of cells which then elongate.

In addition, comparison of apex root cell architecture of the *Atphyl2*.*1-1* mutant and another mutant *Atphyl2*.*1-2* (SALK, with a lower but significant root growth phenotype) to that of the wild type (Col-0) identified several clusters of cell/tissue mis-organization in the mutants at various distances from the root quiescent center. Distances (µm) from the root quiescent center to the transition zone and to the first areas of mis-organized cells were affected in both *Atphyl2*.*1* mutants. Using an immunological approach we further investigated the SNARE Knolle, a specific marker of cell plate formation, and revealed that cell plate formation was significantly affected and is likely responsible for the compromised effectiveness of cell division.

Our results provide a first characterization of the phytolongin AtPHYL2.1 which is involved in root cell plate formation, root cell division and therefore root development.

## Materials and Methods

### Plant lines

The *Arabidopsis thaliana* wild-type ecotype Colombia-0 (Col-0) and two KO mutants were used throughout this study: the mutant *Atphyl2*.*1-1* (RATM53-3274-1) and the mutant *Atphyl2*.*1-2* (SALK_093979). The At*PHYL*2.1 overexpressing line OE3 (p35S:GFP-*PHYL2*.*1*) was crossed with the mutant *Atphyl2*.*1-1* to establish the complemented lines (#12 and #21). *AtPHYL2*.*1* expression level in primary roots of the wild-type ecotype Col-0, the mutant *Atphyl2*.*1-1* (RATM53-3274-1) and the *AtPHYL2*.*1* overexpressing line OE3 (p35S:GFP-*PHYL2*.*1*) are shown in the supplementary Figure S1. All primers used for genotyping and qPCR are listed in Table S1.

### Plasmid preparation and transgenic plants

Sequence data of *AtPHYL2*.*1* can be found in Arabidopsis Information Resource (TAIR) databases under the accession numbers *At4g27840*.

*AtPHYL2*.*1* coding sequences were amplified from Arabidopsis leaf cDNA and cloned by BP reaction in pDONR™221 entry vector (Thermofisher Scientific) using Gateway® cloning technology (Thermofisher Scientific). For overexpression in plants, *AtPHYL2*.*1* entry vector and pK7WGF2 destination vectors were combined by LR recombination using Gateway® cloning technology (Thermofisher Scientific). The newly created vector allows the expression of *GFP-PHYL2*.*1* under the control of the promoter *p35S*. PCR amplification was performed using Q5™ High-Fidelity DNA polymerase at the annealing temperature and extension times recommended by the manufacturer (Biolabs #M04915). PCR fragment and plasmid were respectively purified with NucleoSpin® Gel and PCR cleanup (Macherey-nagel # 740609) and NucleoSpin® Plasmid (Macherey-Nagel #740499). All the entry vectors were sequenced, and sequences were analysed with CLC Mainwork Bench 6. Primers used in this study are listed in table S1. Constructs were transferred into the *Agrobacterium tumefaciens* C58C1 RifR strain harbouring the plasmid pMP90. This strain was used to transform *Arabidopsis thaliana* by floral dip according to Clough et al. (1998).

### RNA extraction, RT–PCR, and quantitative PCR analysis

Plant tissue for RNA extraction, RT–PCR, and qPCR were disrupted using 5 mm stainless steel beads (Qiagen#69989) and Tissuelyser II (Qiagen). Total RNA was extracted from roots 5 days after germination using the RNeasy® Plant Mini kit (Qiagen #74904) according to the manufacturer’s instructions. For full-length transcript analysis, first-strand cDNA was synthesized using SuperScript® II Reverse Transcriptase (ThermoFisher # 18064014) and oligo(dT). Then, mRNAs were treated with DNase I using the DNA-free™ Kit (ThermoFisher # AM1906). Analyses of expression of full-length *AtPHYL2*.*1* by RT–qPCR was performed with the Bio-Rad CFX96 real-time system using GoTaq® qPCR Master mix (Promega # A6002). The transcript abundance in samples was determined using a comparative cycle threshold (Ct) method. The *AtTIP41like* gene (AT4G34270) was used as a constitutively expressed control. The relative abundance of *AtTIP41like* mRNAs in each sample was determined and used to normalize for differences of total RNA level as described in Pascal et al. (2013). For the analysis of the relative abundance of *AtPHYL2*.*1* transcripts at different timepoints (Fig.1), RNA was extracted by mean of the RNeasy® Plant Mini kit (Qiagen #74904) and contaminant genomic DNA was removed by in-column DNase treatment (Qiagen#79254). Then first-strand cDNA was synthesised using SuperScript® IV Reverse transcriptase (ThermoFisher# 18090050) with random hexamers (Promega#C1181). RT-qPCR was performed with a QuantStudio 5 Real-Time PCR System (ThermoFisher) using GoTaq® qPCR Master mix (Promega # A6002). All primers are listed in Table S1.

**Fig 1.**
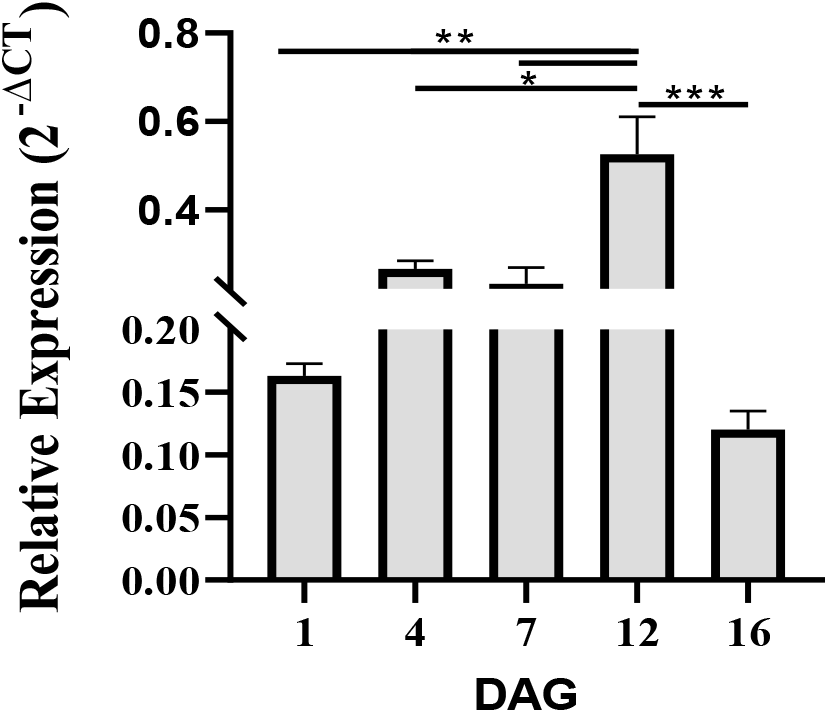
Expression profile of *AtPHYL2*.*1* in *Arabidopsis thaliana* Col-0 seedlings. Expression of *AtPHYL2*.*1* at different time points (DAG = days after germination). Expression levels were analysed using the 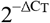 method. *AtACTIN2* (At3g18780) was used as the housekeeping gene. Statistical significance (ordinary one-way ANOVA) is indicated by asterisks: *P < 0.05, **P < 0.005, ***P < 0.001. N = 3 biological replicates, each with 10 seedlings. Error bars = SE (standard error).

### Plant genotyping

Genomic DNA was extracted from leaf material freshly ground in the extraction buffer (25 mM EDTA, 0.25 M NaCl, 0.5% SDS, 0.2 M Tris pH7.5) and centrifuged for 5 min at 20.000 *g*. DNA was precipitated with isopropanol (1:1) and centrifugated 10 min at 20.000 *g*. The DNA pellet was washed with EtOH 70% and resuspended in Tris 10 mM pH8.

Purified DNA was used for genotyping by PCR amplification using GoTaq® Green Master Mix with Standard Buffer (Promega # 7122). All primers are listed in Table S1.

### Phenotypical characterization and growth conditions

Seeds were sterilized by treatment with 95% (v/v) ethanol for 10 s and then repeatedly washed with sterile water and left at 4 °C for 2 days. Seedlings were grown on 1/2 MS agar medium plate (Murashige and Skoog medium 2.2 g/L, sucrose 10 g/L, plant agar 8 g/L, morpholinoethanesulfonic acid (MES) 0.5 g/L, plant agar 8 g/L, buffered to pH 5.8 with KOH) for 5 days (16 h light/8 h darkness). Root length, cell numbers and distances from apex were determined using the ImageJ software (https://imagej.net/).

### Immunocytochemistry, Calcofluor white staining, and confocal laser scanning microscopy

Whole-mount immunolabelling of *Arabidopsis thaliana* roots was performed as described in Boutté and Grebe (2014). In brief, 5-day-old seedlings grown on ½ MS were fixed in 4% paraformaldehyde dissolved in MTSB (50 mM PIPES, 5 mM EGTA, 5 mM MgSO4 pH 7 with KOH) for 1 h at room temperature and washed three times with MTSB. Roots were cut on superfrost slides (Menzel Gläser, Germany) and dried at room temperature. Roots were then permeabilized with 2% Driselase (Merck #D9515), dissolved in MTSB for 30 min at room temperature, rinsed four times with MTSB, and treated for 1 h at room temperature with 10% DMSO+3% Igepal CA-630 (Merck # I3021) dissolved in MTSB. Unspecific sites were blocked with 5% normal donkey serum (NDS; Merck # D9663) in MTSB for 1 h at room temperature. Primary antibodies, in 5% NDS/ MTSB, were incubated overnight at 4 °C and then washed four times with MTSB. Secondary antibodies, in 5% NDS/MTSB, were incubated for 1 h at room temperature and then washed four times with MTSB. Primary antibodies were diluted as follows: rabbit anti-KNOLLE 1:8000 (kind gift of Gerd Jürgens, University of Tübingen, Germany). Dilution of secondary antibody AlexaFluor 555-coupled donkey anti-rabbit IgG (Abcam, ab150062) was 1:800. Calcofluor white 1µg/mL (merck, 18909) was added to stain the cellulose for 20 min, then rinsed twice with MTSB.

Confocal laser scanning microscopy was performed using a Zeiss LSM880 microscope. Laser excitation lines for the different fluorophores were 561 nm for AlexaFluor 555, 405 nm for Calcofluor white. Fluorescence emissions were detected at 570-642 nm for AlexaFluor 555 and 410-524 nm for Calcofluor white. Scanning was performed with a pixel dwell of 3 μs. In multilabelling acquisitions, detection was in sequential line-scanning mode. A Plan-Apochromat 20x/0.8 M27 objective was used. Image analysis was performed using ZEN lite 2.6 2018 (Zeiss) and ImageJ software.

### Statistical analysis

All data analysed were unpaired (samples independent from each other). Normal distribution (Gaussian distribution) of the data sets was tested using Shapiro–Wilk normality test. Our data sets were not normally distributed so we performed non-parametric tests. We also used non-parametric tests on data sets for which n <10. To compare multiple data sets, Kruskal– Wallis tests were used as non-parametric tests. To identify statistically significant differences between two data sets, we used Mann–Whitney test as non-parametric test. All statistical analyses were performed with the R i386 3.1.0 software. P-values were as follows: NSP-value>0.01 (not significant), *P<0.05, **P<0.01 and ***P<0.001.

### Cell length measurements and ImageJ analysis

Immediately before acquisition, root apical meristems and transition zones from 5 day-old seedlings were either stained with Propidium Iodide (ThermoFisher #P21493) (10mg/ml, in ½ MS) or with 2% propidium iodide solution (Merck#P4170) in the mounting media. Images were acquired using either a Zeiss LSM880 confocal microscope or a Leica SP8 Stellaris confocal microscope using a 40× objective (dry). The dye was excited at 541 nm, fluorescence was detected at 560-690 nm. Cells were measured in the epidermis, from the first epidermal cell until the last epidermal cell visible on the images, measuring points being the centre of each cell wall. The TZ was positioned at the level of the last isodiametric cell before the first elongating cell in the cortex tissue. Images were processed using a homemade ImageJ macro program (the whole program can be viewed in the supplementary data) that provides a segmented, binary image for cell length measurements.

## Results

### Primary root length is significantly reduced in the Atphyl2.1-1 loss-of-function mutant

To investigate the potential role of PHYL2.1 in *Arabidopsis thaliana* development and to compare loss-of-function mutants of *PHYL2*.*1* with the wild type in the most accurate manner, we first determined the relative expression level of *PHYL2*.*1* up to 16 days after germination. As indicated in Fig.1, the level of expression of *PHYL2*.*1* is higher and relatively stable between 4 and 12 days after germination. Since the root meristem size and the transition zone are established as early as 5 days after germination (de Nittis et al., 2025), and that we wanted to look at the role of *PHYL2*.*1* as early as possible in root growth, we decided to grow roots at 5 days after germination and to carry out our analysis at this timepoint.

*Arabidopsis thaliana* wild type ecotype Colombia-0 (Col-0; WT) and the loss-of-function mutant *Atphyl2*.*1-1* (RATM53-3274-1) were grown vertically in 16 h light/8 h darkness for 5 days and root length of seedlings was measured using the ImageJ software. Primary root length (Fig.2) was significantly reduced in the *Atphyl2*.*1-1* mutant compared to the WT seedlings. The mean length of the roots in the *Atphyl2*.*1-1* mutant did not exceed 20-25% of the length of the roots in the WT seedlings, indicating a strong disturbance of primary root growth in the *Atphyl2*.*1-1* mutant, in comparison to WT.

**Fig 2.**
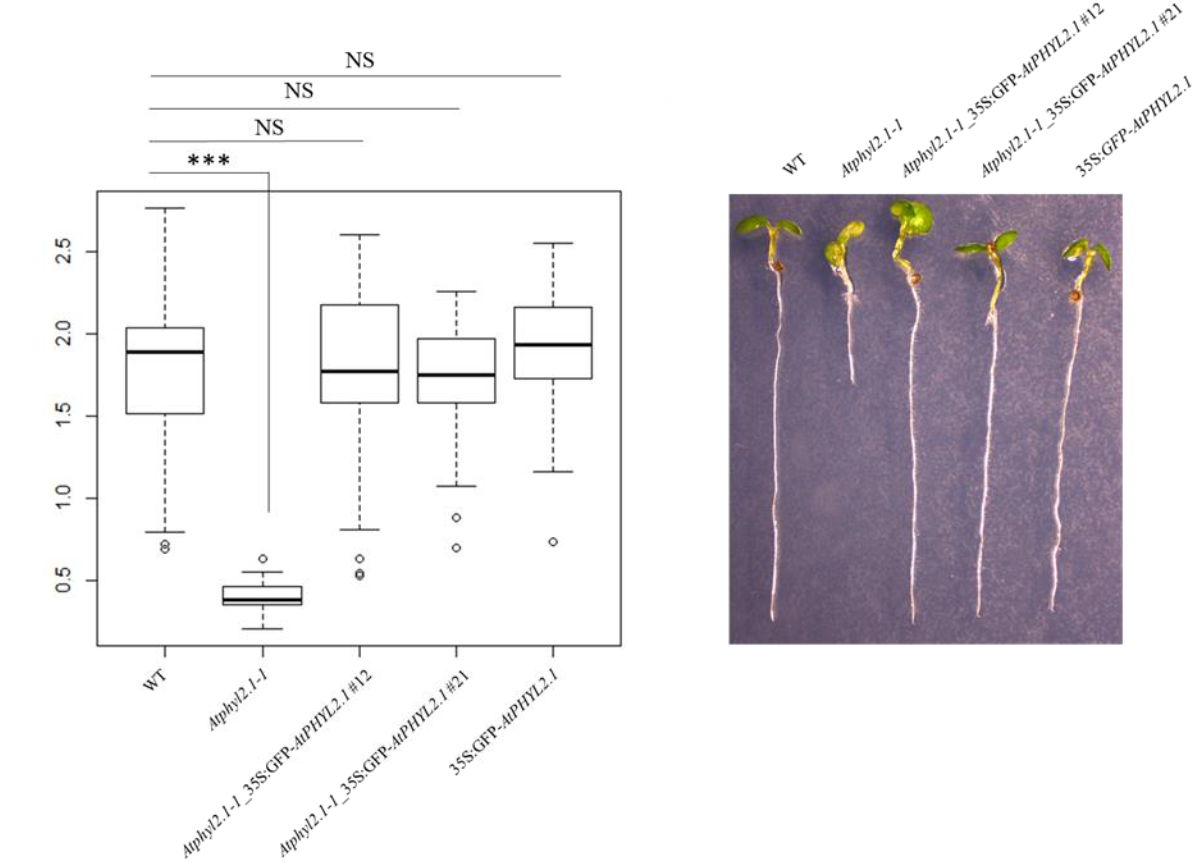
Primary root length is significantly reduced in the *Atphyl2*.*1-1* mutant and rescued by complementation with *AtPHYL2*.*1* overexpression. Primary root length of wild type (WT), Atphyl2.1-1 mutant, 35S:GFP-AtPHYL2.1 overexpression line 3 and two complemented lines (Atphyl2.1-1_35S:GFP-AtPHYL2.1 #12 and #21) were measured 5 days after germination. NSP-value>0.01 (not significant), ***P<0.001 (p-value < 2.2e-16); n=68.

To confirm that loss of AtPHYL2.1 was responsible for the observed primary root length phenotype, we crossed an overexpression line OE3 of *AtPHYL2*.*1* (35S:GFP-*AtPHYL2*.*1*, supplementary Fig.S1) with the mutant line *Atphyl2*.*1-1* and again measured seedling root length. Primary root length was recovered in two independent complementation lines (*Atphyl2*.*1-1*_35S:GFP-*AtPHYL2*.*1* #12 and #21, Fig.2) with mean root lengths being comparable with WT seedlings, the overexpression line used for the complementation and the rescued lines of *Atphyl2*.*1-1* mutant. The recovery of primary root length in the complemented lines indicates that the phytolongin AtPHYL2.1 is involved in primary root growth.

### Cell division but not cell elongation appears to be affected in the Atphyl2.1-1 mutant

Since primary root length was significantly reduced in the *Atphyl2*.*1-1* mutant, we wondered whether cell division and/or cell elongation were also affected in this line. To investigate these possibilities, we developed a tool to measure the number of cells and their size from the quiescent centre to the cell division zone and part of the cell elongation zone (Fig.3).

**Fig 3.**
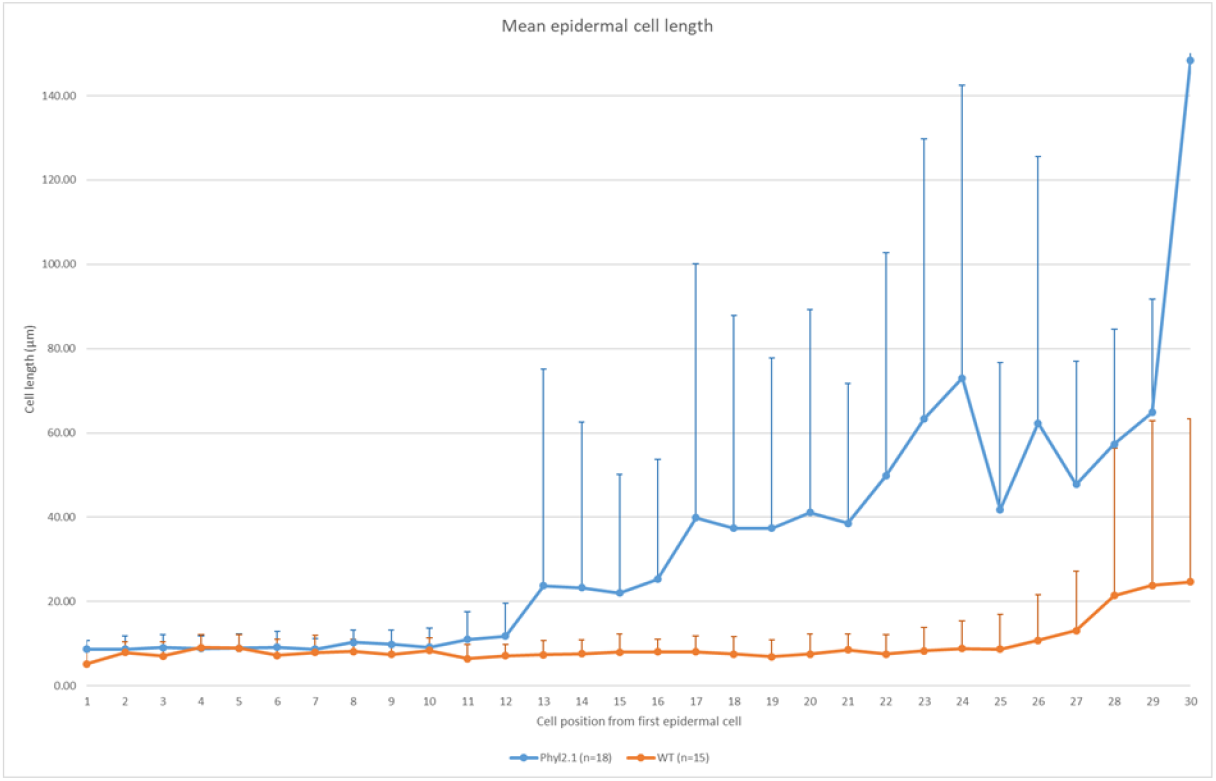
Cell division but not cell elongation is reduced in the *Atphyl2*.*1-1* mutant. Cell size was measured using a homemade ImageJ macro program (shown in supplementary data). The graph indicates the cell length of the first 30 epidermal cells from the quiescent centre of the primary roots of WT (n=15) and *Atphyl2*.*1-1*. (n=18).

Cell length started to increase only at the level of cell 28, from the quiescent centre, in the WT roots whereas cell length started to increase already at the level of cell 13 in the *Atphyl2*.*1-1* mutant (Fig.3). Therefore, we observed that the cell division zone in the WT roots extended up to ∼300µm from the quiescent centre and that cellular elongation started only above this distance. However, in the *Atphyl2*.*1-1* mutant the cell division zone was restricted to ∼150-200 µm from the quiescent centre and therefore elongation started closer to the quiescent centre.

We determined the mean number of cells at about 200 µm of the quiescent centre to be equal to 26 and 19, for the WT and *Atphyl2*.*1-1* mutant, respectively. At 900µm and 500µm respectively from the root quiescent centre, the final cell size of WT and *Atphyl2*.*1-1* is comparable. However, the mean number of cells overall was only ∼24 cells/900µm for the *Atphyl2*.*1-1* mutants analysed compared to ∼58 cells/900µm for the WT seedlings.

Since root cells from WT and *Atphyl2*.*1-1* mutant reached the same size in the elongation zone, all these data indicate clearly that the cell division rather than the cell elongation was strongly affected in the *Atphyl2*.*1-1* mutant. Therefore, the decrease of root length in the *Atphyl2*.*1-1* mutant could be at least partly explained by a lower number of cells (through a lower efficiency of cell division) which could then elongate to enable root growth.

### Cell/tissue mis-organization in the Atphyl2.1-1 mutant

Given the observed perturbations to cell division in the *Atphyl2*.*1-1* mutant we next analysed the cell architecture in the root apex of the mutant in comparison to that of the WT. In this approach we also used another mutant from SALK collection (*Atphyl2*.*1-2)* which also showed a significant root growth phenotype but to a lesser extent compared to *Atphyl2*.*1-1* mutant (Fig.4B). It was interesting to investigate cell misorganization in the root apex at two different levels of root growth disturbance with the two different mutants.

**Fig 4.**
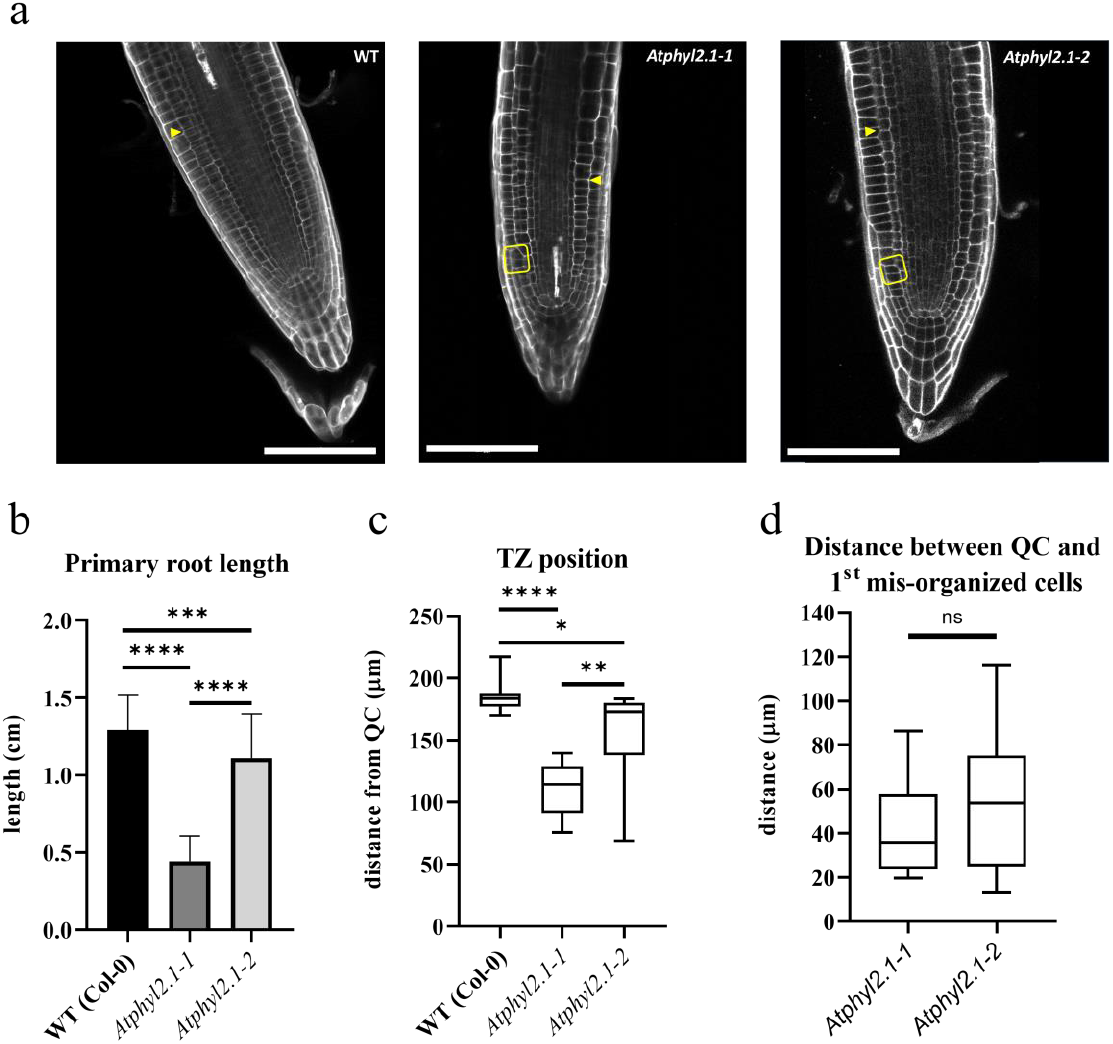
Perturbations in cell division and transition zone positioning in *Atphyl2*.*1-1* and *Atphyl2*.*1-2* mutants. (a) Apex root cell architecture of *Atphyl2*.*1-1* and *Atphyl2*.*1-2* mutants compared with wild type (WT), visualized by confocal microscopy after propidium iodide staining of cell walls. Yellow arrowheads indicate the transition zone (TZ). Yellow boxes indicate representative regions of mis-organized cells in mutants compared with corresponding regions in WT roots. Scale bar = 100 µm. (b) Primary root length of WT, *Atphyl2*.*1-1*, and *Atphyl2*.*1-2* mutants measured 5 d after germination (N >39). (c) Distance from the quiescent centre to the root TZ in WT and mutants. (d) Distance from the quiescent centre to the first region of mis-organized cells in mutants (n > 8). Most clusters of highly mis-organized cells occurred early, at 20–60 µm from the quiescent centre. Statistical significance (ordinary one-way ANOVA) indicated by asterisks: *P < 0.05, **P < 0.01, ***P < 0.005, ****P < 0.001. Error bars = SD.

Mis-organized areas in the mutants are highlighted in yellow boxes (Fig.4A) as examples of the strong disturbance of tissue organization which is observed in the *Atphyl2*.*1* mutants as compared to similar areas in the WT line. Distances (µm) from the root quiescent centre to the transition zone (Fig.4C) were shortened in both mutants but not to the same extent, and distances from the root quiescent centre to the first areas of mis-organized cells (Fig.4D) were similar in both *Atphyl2*.*1* mutants. Although the first clusters of highly mis-organized cells could be observed at various distances from the quiescent centre in the roots of both mutants analyzed, it is clear that cell division and cell organization disturbances occurred quite early.

### Cell plate formation is disturbed in the Atphyl2.1-1 mutant

In higher plant cytokinesis, the new plasma membrane and cell wall originate by vesicle fusion in the plane of cell division. Membrane vesicles delivered to the cell-division plane fuse to form the partitioning membrane, which, in *Arabidopsis thaliana*, requires the cytokinesis-specific Qa-SNARE KNOLLE (Lauber et al; 1997; Park et al. 2018). Consequently, this syntaxin can be considered as a very reliable marker to study cell division in root cells. Therefore, in order to determine whether cell plate formation was disturbed in the *Atphyl2*.*1-1* mutant, we performed an immunolocalization of KNOLLE using a whole-mount immunolabelling approach on *Arabidopsis thaliana* roots as described in Boutté and Grebe (2014) with calcofluor labelling on the cell wall. Using this approach, we were able to count KNOLLE-labelled cell plates on different depth sections in both *Atphyl2*.*1-1* mutant and WT roots and determine the ratio of the number of cell plates per µm of depth for the two conditions (Fig.5). We observed a significant, ∼3-fold decrease of KNOLLE-labelled cell plates in the *Atphyl2*.*1-1* mutant in comparison to WT. This decrease was corrected in the complementation line with even an increase of the number of cell plates labelled. The decrease of KNOLLE-labelled cell plates in the *Atphyl2*.*1-1* mutant strongly suggests an important disturbance of root cell division in the loss-of-function mutant.

**Fig 5.**
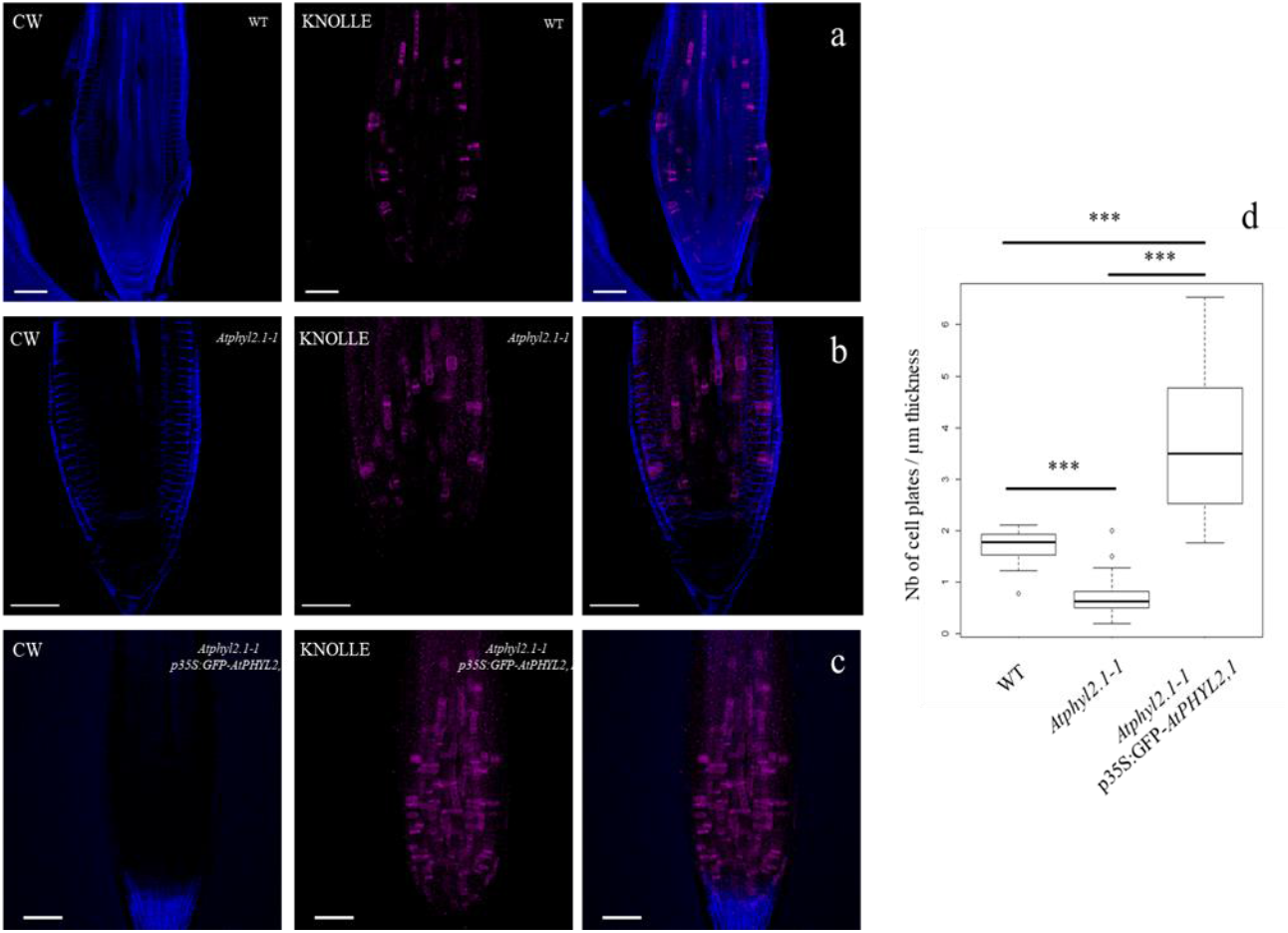
Cell plate formation is significantly reduced in the *Atphyl2*.*1-1* mutant. Immunolocalization of the cell plate marker KNOLLE in root cells from Arabidopsis wild-type (WT) (a), *Atphyl2*.*1-1* mutant (b) and *Atphyl2*.*1-1*_35S:GFP-*AtPHYL2*.*1* #21 (c). Cell wall (CW) is labelled by calcofluor. KNOLLE dependent cell plates were counted at different depths in complementation line, mutant and wild type sections and the ratio of the number of cell plates per µm of depth were determined (d). Scale bars represent 50 µm. n = 18 roots for the WT and n = 38 roots for the mutant, and n = 22 roots for the complementation line. Statistics were done by non-parametric Wilcoxon rank sum test, ***P-value < 0.001 (WT vs At*phyl2*.*1-1* p-value = 3.263e-08 ; WT vs complementation line p-value = 1.056e-06; *Atphyl2*.*1-1* vs complementation line p-value = 1.837e-10). ***P-value < 0.001.

In addition to the decrease in cell division efficiency, we also observed some abnormal formation of cell plates in the *Atphyl2*.*1-1* mutant (Fig.6). Abnormal cell plates were largely observed in the areas of clusters of mis-organized cells closer to the quiescent center (Fig.6), indicating that the mis-organisation of cells and disturbance of cell division occurred very early in root growth for the *Atphyl2*.*1-1* mutant.

**Fig 6.**
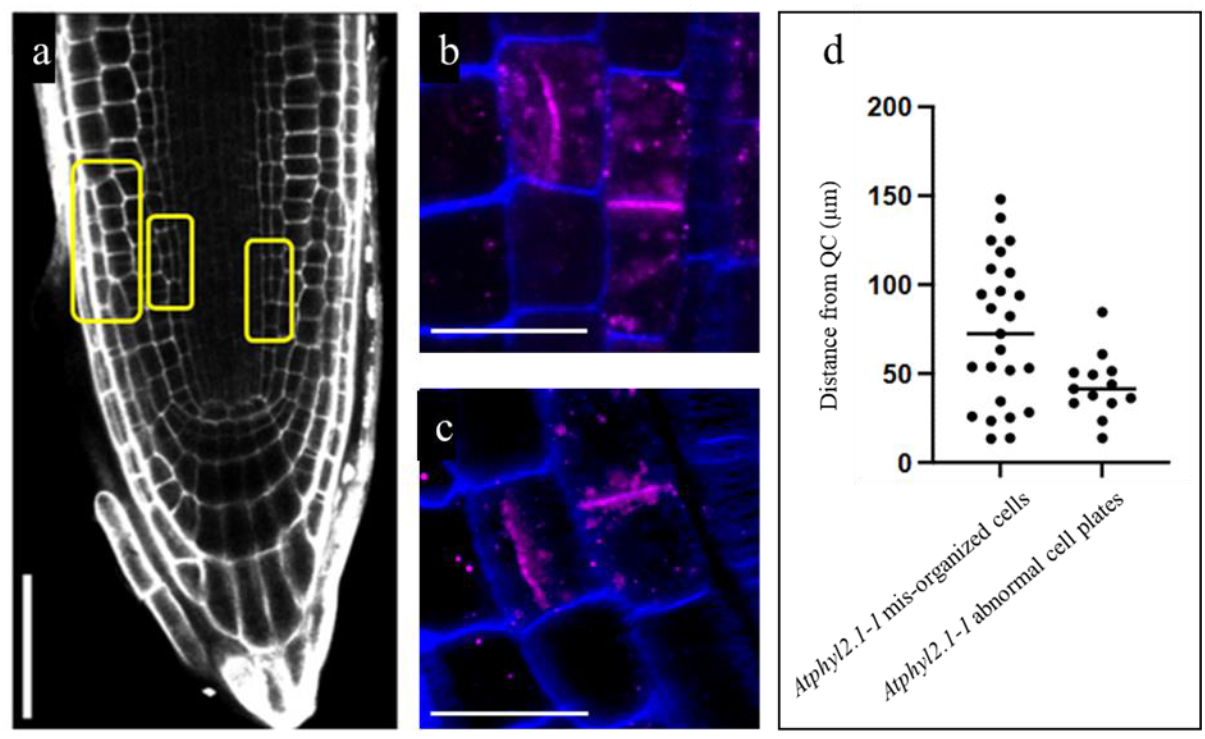
Abnormal cell plates are observed in the first areas of clusters of mis-organized cells in *Atphyl2*.*1-1* mutant. (a) Cell wall staining of *Atphyl2*.*1-1* mutant apex root with propidium iodide. Yellow boxes highlight representative examples of mis-organized areas. Scale bars 50 µm. Distances from the root quiescent center to the first areas of mis-organized cells in the *Atphyl2*.*1-1* mutant were measured (n=13). The majority of the clusters of highly mis-organized cells were observed very early at distances from the quiescent center comprised between 20 and 60 µm. (b,c) Immunolocalization of the cell plate marker KNOLLE in root cells from *Atphyl2*.*1-1* mutant illustrates atypical cell plates. 25 abnormal cell plates were observed among 13 of the 39 roots analyzed. Cell wall (CW) is labelled by calcofluor. Scale bars 10 µm. (d) Distances from the root quiescent center of the abnormal cell plates were compared to those of the areas where mis-organized cells were observed in *Atphyl2*.*1-1*. Measurements were realized with imageJ.

Taken together, these data indicate a critical disturbance of cell division and tissue organization in the loss-of-function *Atphyl2*.*1-1* mutant which, in turn, is likely responsible for the observed reduced root growth.

## Discussion

Phytolongins represent a relatively newly described set of proteins from the longin family proteins that have been diversified in plants (Vedovato et al., 2009). The longin domain is a building block of the eukaryotic trafficking machinery and hence it is shared by proteins from the Adaptor Protein (AP) clathrin adaptor, TRAnsport Protein Particle (TRAPP), Signal Recognition Particle (SRP) and COPI complexes (De Franceschi et al. 2014). Therefore, this domain can be associated with several different protein domains/regions, and in the *sensu stricto* longins subfamily (including VAMP7, Sec22 and Ykt6), it is associated with the SNARE motif. Notably, although based on homology, phytolongins belong to the *sensu stricto* longins subfamily and share with them the conserved longin domain, they do not possess a SNARE motif but do contain instead another uncharacterized “Phyl region” (Vedovato et al., 2009; De Marcos Lousa et al, 2016). Phytolongins are not the only *sensu stricto* longins that miss a SNARE motif, as this domain is absent also in mammalian Sec22b homologues Sec22a/c; however, both types of non-SNARE longins are still involved in subcellular trafficking (Rossi et al., 2004; De Marcos Lousa et al, 2016). Considering the role that the SNARE motif generally plays in subcellular trafficking, its absence in phytolongins might suggest these proteins are unable to mediate fusion events, while possibly regulating trafficking pathways specific to land plants physiology and/or development.

It was therefore interesting to address this “functional” point and for that, we investigated the role of the phytolongin PHYL2.1 in primary root growth after having determined a strong root growth phenotype in an *Atphyl2*.*1* mutant (Fig.2). Primary root length was dramatically reduced in the *Atphyl2*.*1-1* mutant (mean root length of *Atphyl2*.*1-1* were ∼20-25% of that of wild type Col-0). To determine if the loss of *AtPHYL2*.*1* was responsible for the observed root phenotype, we complemented the mutant line *Atphyl2*.*1-1* by crossing with an overexpression line. Two independent complementation lines (*Atphyl2*.*1-1* 35S:GFP-*AtPHYL2*.*1* #12 and #21) clearly showed a recovery of primary root length (Fig.2). Therefore, this recovery suggests that AtPHYL2.1 is involved in primary root growth.

We then wondered whether the observed perturbation in root growth in the *Atphyl2*.*1-1* mutant was underpinned by alterations in cell elongation and/or cell division. Comparison of the characteristics of the roots of the *Atphyl2*.*1-1* mutant with those of the WT led to the following conclusions: *i*) the cell division zone was significantly reduced (∼150-200µm) in the *Atphyl2*.*1-1* mutant in comparison to the WT (∼300µm); *ii*) cell elongation started closer to the quiescent centre in the *Atphyl2*.*1-1* mutant than in the WT; *iii*) the mean number of cells in the division zone was significantly reduced in the *Atphyl2*.*1-1* mutant (mean 19 cells/200µm) in comparison to the WT (mean 26 cells/200µm); *iv*) whilst root cells from both WT and *Atphyl2*.*1-1* mutant eventually reached comparables sizes in the elongation zone, this occurred at ∼800-900µm from the quiescent centre in the WT but at ∼400-500µm from the quiescent centre in the *Atphyl2*.*1-1* mutant (Fig.3 and Fig.4). Taken together, these findings indicate that impaired cell division, rather than compromised cell elongation, constitutes the primary mechanism underlying the observed reduction in root length in the *Atphyl2*.*1-1* mutant, with subsequent cellular elongation serving as a compensatory mechanism to maintain root growth.

In order to determine whether cell plate formation was disturbed in the *Atphyl2*.*1-1* mutant, we performed an immunolocalization of the well characterised cytokinesis-specific marker, KNOLLE (Lauber et al; 1997; Park et al. 2018). A high disbturbance of KNOLLE-labelled cell plates was found in the *Atphyl2*.*1-1* mutant (Fig.5a-c) and the ratio of the number of cell plates/µm of depth was strongly decreased in the mutant (Fig.5d) which was corrected in the complementation line. This clearly indicates that the drop in the density of KNOLLE-labelled cell plates in the *Atphyl2*.*1-1* mutant is linked somehow to the function of AtPHYL2.1. We also observed some abnormal formation of cell plates in the *Atphyl2*.*1-1* mutant (Fig.6b-d), together with an early mis-organization of the tissues in the roots of the *Atphyl2*.*1-1* mutant (Fig.6a).

Therefore, all these results combined demonstrate that a critical disturbance of cell division and tissue organization appears in the *Atphyl2*.*1-1* mutant which readily explains the strong deficiency of root growth that is observed in the *Atphyl2*.*1-1* mutant.

What could be the function of AtPHYL2.1, a phytolongin, in *Arabidopsis* root cell division and tissue organization? The lack of a SNARE domain does not favour a function of AtPHYL2.1 in membrane fusion and therefore a function in vesicle trafficking to the cell plate seems unlikely. However, phytolongins were found to be characterized by the presence of a NIE hydrophilic loop in the Phyl domain (replacing the SNARE domain), but similar to that in the SNARE motif of Sec22 (Vedovato et al., 2009). As a consequence the question of AtPHYL2.1 function in membrane trafficking still persists. As discussed in de Marcos Lousa et al. (2016), the presence of this hydrophilic “NIE-like” loop in the Phyl domain of AtPHYL2.1 could suggest that the longin domain of AtPHYL2.1, by interacting with the “NIE-like” loop of the Phyl domain, could fold in a “closed” conformation incompatible with the formation of a SNARE complex and therefore membrane fusion.

The lack of a classical SNARE motif in AtPHYL2.1 leaves open the possibility that the “NIE-like” loop could also interact with the longin domain of other SNAREs to regulate their availability to form SNARE complexes. In addition, the interaction of the “NIE-like” loop of the Phyl domain of AtPHYL2.1 could also interact with the longin domain of AtPHYL2.1, adding another level of regulation.

Finally, according to possibilities already mentioned in de Marcos Lousa et al. (2016), the localisation of AtPHYL2.1 to the ER may mean that AtPHYL2.1 interacts with specific reticulon and/or reticulon-like proteins, major ER morphogenic proteins (Kriechbaumer et al., 2015 and references herein). Such interactions with proteins of the reticulon family may have a role in ER organization and/or at the ER-PM interface, further implicating AtPHYL2.1 in the efficiency of cell plate formation and regulation of tissue organization.

## Supporting information

Table S1, Fig. S1 and macro program

## Supplementary data

Supplementary Table S1 shows the list of the primers used in this study. Supplementary Fig.S1 shows the expression level of At*PHYL2*.*1* in primary roots. Supplementary Materials and Methods : the Macro Fiji program is presented.

## Acknowledgments

Imaging was done at the Bordeaux Imaging Center (UAR CNRS-University of Bordeaux 3420, http://www.bic.u-bordeaux2.fr), member of the France-BioImaging national infrastructure (http://france-bioimaging.org/). We thank Leslie Bancel-Vallée, Guillaume Bouyssou and Chiara Banzato for their involvement in some of the experiments. We thank Magali Grison for her help with the immunolabeling protocol.

## Author contributions

VW-B, LB, MB, EB, MZ, EF, FF and PM: conceptualization; VW-B, MB, FD-D, LM-P, YLN and CM performed the experiments; EB performed the overexpressed lines which were used by VW-B to cross with the mutant lines to create the complemented lines, and had significant input into the editing of the final manuscript; all authors analyzed and discussed the data; PM: supervision, writing the article with contributions of all the authors; VW-B, LB and PM are responsible for contact and communication.

## Conflict of interest

The authors declare no conflict of interest.

## Funding

This work was supported by CNRS and the University of Bordeaux which PM thanks for their continuous support all along his career.

## Data availability

The data supporting the findings of this study are available within the paper and within its supplementary data.

## Notes

### Competing Interest Statement

The authors have declared no competing interest.

