## Supplementary material for "The phytolongin AtPhyl2.1 is involved in cell plate formation and root development": Table S1, Fig. S1 and macro program

**Supplementary data**

**Supplementary Table S1 : List of primers used in this study**

| GENE NAME | AIM | PRIMER SEQUENCE (5'-3') |
|---|---|---|
| *AtPHYL2.1* | Cloning | Forward_ggggacaagtttgtacaaaaaagcaggctggATGACTTCGAATCAACGTATG |
| | | Reverse_ggggaccactttgtacaagaaagctgggtgTTACCCATCGATGCATTGAAAC |
| *AtPHYL2.1* | RT-qPCR Fig.1 | Forward_ GGAAGACGGATCTACTGCGG |
| | | Reverse_ TCCCTCGCTATACTCCTCGG |
| *AtACTIN2* | RT-qPCR Fig.1 | Forward_AAGCTCTCCTTTGTTGCTGTT |
| | | Reverse_GACTTCTGGGCATCTGAATCT |
| *AtPHYL2.1* | RT-qPCR Fig.S1 | Forward_ACTCCTTTACCTCCTCCTCC |
| | | Reverse_CGCTGAGATTGCTGCTATTG |
| *AtTIP41like* | RT-qPCR Fig.S1 | Forward_GCTCATCGGTACGCTCTTTT |
| | | Reverse_TCCATCAGTCAGAGGCTTCC |
| *Atphyl2.1-1* (RATM53-3274-1) | Genotyping | Forward_TTTCGCTGAGATTGCTGCTA |
| | | Reverse_ATCTTGCGGACAAAAATGCT |
| | | TDNA_CCGTCCCGCAAGTTAAATATG |
| *Atphyl2.1-2* (SALK_093979) | Genotyping | Forward_TACCTTGGTGAAAGTGCCCT |
| | | Reverse_TTGTGGCGTACATGACCTGA |

| | | |
|---|---|---|
| | | TDNA_Forward ATTTTGCCGATTTCGGAAC |

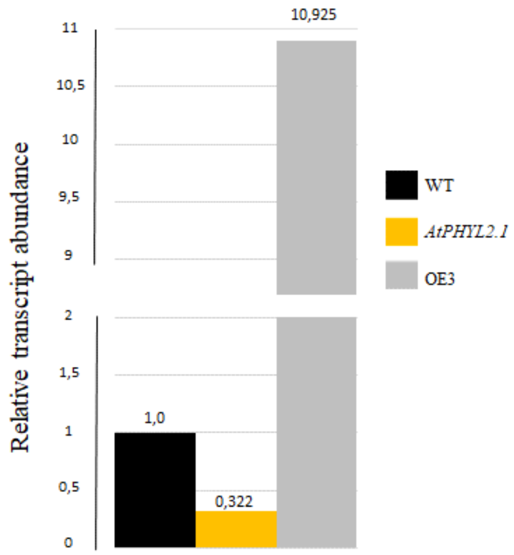


**Supplementary Fig.S1: At*PHYL2.1* expression level in primary roots**
The results were analyzed using the 2-ΔCT method. *AtTIP41like* (At4g34270) has been used as a reference gene. (A) Expression level of At*PHYL2.1* gene in primary roots from 5 day-old seedlings.

### Supplementary Materials and Methods : Fiji macro program

```
//Macro written by Matthieu Buridan
//Variables declaration

//Directories
var inDir = "";
var outDir = "";
var files = "";

//Image names
var ori = "";
var img1 = "";
var img2 = "";
var DoG = "";
var binary = "";
var oriSave = ""

//Processing
var sigma1 = "";
```

```
var sigma2 = "";

var nbSlices ="";
var suffix = "";

var imgType = "";


//------------------------------------------------------------------------

//Adds the single file macro to the toolset

macro "Single_file Action Tool -
N66C000C111C222D52C222C333D8cD9cDacDbcDccDdcDecC333D62D72D82D92Da2Db2Dc2Dd2
De2C333D8aD9aDaaDbaDcaDdaDeaC333C444C555D88D98Da8Db8C555D43D44D45D46D47D48D
49D4aD4bD4cD4dD4eC555C666D42C666Dc8C666C777D7cC777D87D97Da7Db7C777D7aC777C8
88Dc7C888D53C888D78C888D8eD9eDaeDbeDceDdeDeeC999D77C999D54D55D56D57D58D59D5
aD5bD5cD5dD5eC999CaaaD8dD9dDadDbdDcdDddDedCaaaCbbbD7eCbbbD7dCcccD89D99Da9Db
9Dc9CcccDd9CcccDe9CdddD51D61D71D81D91Da1Db1Dc1Dd1De1CdddD8bD9bDabDbbDcbDdbD
ebCdddD79CdddCeeeD41CeeeD7bCeeeD63D73D83D93Da3Db3Dc3Dd3De3CeeeDd8CeeeDd7Cee
eCfffD6cCfffD33D34D35D36D37D38D39D3aD3bD3cD3dD3eD68D6aCfffD32D67D6eCfffD6dC
fffD31D69D6bBf0C000D24D25D35D45C000D34C000D23C000D55C000C111D12C222D44C222D
13C222C333D14D15C333D0cD1cD2cC333D02C333D66C333D0aD1aD2aC333C444D33D67D68D6
9D6aD6bD6cD6dD6eC444C555D56C555D3cC555C666D3aC666D26D36D46C666C777D22C777D6
5C777C888D16C888D0eD1eD2eC999CaaaD54CaaaD3eCaaaD0dD1dD2dCaaaCbbbD3dCbbbCccc
D57D58D59D5aD5bD5cD5dD5eCcccD43CcccD09D19D29CdddD01CdddD03CdddD11D39CdddD0b
D1bD2bCdddD3bCeeeD32CeeeD04D05CeeeCfffD4aD4cD76D77D78D79D7aD7bD7cD7dD7eCfff
D06CfffD64CfffD21D4eCfffD4dD75CfffD49CfffD4bCfffD53CfffB0fC000C111C222D58C2
22D02D12D22C222D08D18D28D38D48C222C333D68C333C444D67C444D60D61D62D63D64D65D
66C444D32C555C666D00D10D20C666C777C888D30C888C999CaaaD57CaaaCbbbD09D19D29D3
9D49D59CbbbCcccD50D51D52D53D54D55D56CcccD01D11D21CcccD69CcccD31CcccCdddD03D
13D23CdddCeeeD33CeeeD07D17D27D37D47CeeeD42CfffD70D71D72D73D74D75D76D77CfffD
78CfffD40CfffD41CfffD79CfffD43CfffNf0C000C111D58C111C222D82D92Da2Db2Dc2Dd2D
e2C222D68D78D88D98Da8Db8Dc8Dd8De8C222C333C444C555D40D41D42D43D44D45D46D47C5
55D48C555C666D80D90Da0Db0Dc0Dd0De0C666D72C666C777C888D57C888C999D70C999D50D
51D52D53D54D55D56C999CaaaCbbbD59D69D79D89D99Da9Db9Dc9Dd9De9CbbbCcccD81D91Da
1Db1Dc1Dd1De1CcccD49CdddD71CdddD83D93Da3Db3Dc3Dd3De3CdddCeeeD67D73D77D87D97
Da7Db7Dc7Dd7De7CeeeCfffD62CfffD30D31D32D33D34D35D36D37CfffD38D60CfffD39D61D
63"{
	setBatchMode("show"); //Desactivate batch mode as manual processes
must be conducted
	oneFile();
}

//------------------------------------------------------------------------

//Adds the multiple file macro to the toolset

macro "Multiple_files Action Tool -
N66C000D60C000D70D80D90Da0Db0Dc0Dd0De0C000C111C222D34C222D7eD8eD9eDaeDbeDce
DdeDeeC222C333D6eC333D35C333D36D37D38D39D3aD3bD3cD3dD3eDe4C333D79D89D99Da9C
333C444D63C444D44C444D69C444D61C555D54D64C555D53D74D84D94Da4Db4Dc4Dd4C555D7
cD8cD9cDacDbcDccDdcDecC555C666D6cC666D62C666C777C888Db9C888D50D51D52C888C99
9D7aD8aD9aDaaC999D7bD8bD9bDabDbbDcbDdbDebC999CaaaD6aCaaaD43D73D83D93Da3Db3D
c3Dd3De3CaaaD6bCaaaD33CaaaCbbbDe5CbbbDe6CbbbDe7CbbbD71D81D91Da1Db1Dc1Dd1De1
CbbbCcccDbaCcccD7dD8dD9dDadDbdDcdDddDedCcccD6dCcccCdddD25D26D27D28D29D2aD2b
D2cD2dD2eD5eCdddD24CdddD45D59CdddCeeeD46D47D48D49D4aD4bD4cD4dD4eCeeeD5cCeee
De8CeeeD5aCeeeD5bCeeeCfffD23CfffD78D88D98Da8CfffD68CfffD5dCfffDb8CfffD55D65
D75D85D95Da5Db5Dc5Dd5CfffD58CfffDc9Bf0C000D16D32D33D42D43D53C000D26C000D31C
000D17D27D37C000D15C000D05D06C000D00D07D10D20C000D63C000C111D47C111D04C111C
222D52C222D36C222D74C222D0eD1eC222C333D75D76D77D78D79D7aD7bD7cD7dD7eC333C44
```

```
4D41C444D21C444D25D48C444D64C444C555D34C555D44D54C555D22D23C555D0cD1cC555C6
66D30C666D49D4aD4bD4cD4dD4eC666C777D73C777D14C777D24C777C888D58D59D5aD5bD5c
D5dD5eC888C999D2eC999D0bD1bC999D62C999CaaaD46CaaaD57CaaaD18D28D38CaaaD08Caa
aD2cCaaaCbbbD03CbbbD11CbbbD01CbbbCcccD51CcccD35CcccD0dD1dCcccD2bCcccCdddD85
D86D87D88D89D8aD8bD8cD8dD8eCdddD84CdddD65CdddCeeeD66D67D68D69D6aD6bD6cD6dD6
eCeeeD2dD40CeeeCfffD13D83CfffD72CfffD56CfffD45CfffD12D55CfffD61CfffB0fC000D
4aC000D0aD1aD2aD3aC000C111C222D76C222D56C222C333D75C333D70D71D72D73D74C333C
444D04D14C444D66C444D49C555D01D11C555D57C555C666D40D41D42D43D44D45D46D47D48
C666C777C888D55C888D03D13D50D51D52D53D54D58D59D5aC888C999CaaaD24D67CaaaD02D
12CaaaD21CaaaD77CaaaCbbbD09D19D29D39CbbbCcccD23CcccD00D10CdddD22CdddD80D81D
82D83D84D85CdddD86CdddD65CdddCeeeD60D61D62D63D64CeeeD20CeeeCfffD87CfffD05D1
5CfffD25CfffNf0C000D4aC000D5aD6aD7aD8aD9aDaaDbaDcaDdaDeaC000C111D3aC111C222
C333D39C333D30D31D32D33D34D35D36D37D38C333C444D74D84D94Da4Db4Dc4Dd4De4C444C
555D64D71D81D91Da1Db1Dc1Dd1De1C555C666D61C666C777C888D73D83D93Da3Db3Dc3Dd3D
e3C888C999D63C999CaaaD72D82D92Da2Db2Dc2Dd2De2CaaaD49CaaaD62CaaaCbbbD59D69D7
9D89D99Da9Db9Dc9Dd9De9CbbbCcccD70D80D90Da0Db0Dc0Dd0De0CdddD60CdddD20D21D22D
23D24D25D26D27D28D29D2aCdddD54CdddCeeeD51CeeeD40D41D42D43D44D45D46D47D48Cee
eD53CeeeD52CfffD50CfffD75D85D95Da5Db5Dc5Dd5De5CfffD65CfffD55"{
	setBatchMode("show"); //Desctivate batch mode as manual processes
must be conducted
	multipleFiles();
}

//--------------------------------------------------------------------------

//Starts the process in single file mode

function oneFile(){
	GUI(false); //Calls the GUI function in single file mode
	processSingleFile(); //Calls the main process function
}

//--------------------------------------------------------------------------


function multipleFiles(){
	GUI(true); //Calls the GUI function in multiple file mode
	files=getFileList(inDir);
	//Scans the file list and open files with .czi extension only
	for(i=0; i<files.length; i++){
		if(endsWith(files[i], suffix)){ //Suffix is .czi
			openFile(inDir, files[i]); //Call the open file
function
			nbSlices=nSlices;
			if (nbSlices==1){
				processSingleFile(); //Starts the processing
function
			}
		}
	}
	waitForUser("Done !");
}

//--------------------------------------------------------------------------


function processSingleFile(){
	//Function that calls every other function for the processing
	checkFormat();
	preProcessing();
	differenceOfGaussians();
```

```
        processingSegmentation();
        manualSegmentation();
        measurement();
        saveData();
}

//-------------------------------------------------------------------------


//Calls Bioformat with the path of the image to open
function openFile(inDir, file){
        inputfile = inDir + File.separator + file; //Concatenate the path
and the filename into inputfile variable
        run("Bio-Formats", "open=["+ inputfile +"] autoscale
color_mode=Default rois_import=[ROI manager] view=Hyperstack
stack_order=XYCZT"); //Open the .czi file with bioformats
}

//-------------------------------------------------------------------------


//GUI that asks for folder path and sigma parameters for the difference of
gaussians
function GUI(batch){

        Dialog.create("Parameters");

        //Directories parameters, input is activated if batch is true.
output is mandatory
        Dialog.addMessage("Saving options", 18)
        if(batch){
                Dialog.addDirectory("Input_folder", inDir);
        }
        Dialog.addDirectory("Output_folder", outDir);
        Dialog.addString("Format", ".tif");

        Dialog.addMessage("Difference of Gaussians", 18);
        Dialog.addSlider("Sigma 1", 1, 5, 2);
        Dialog.addToSameRow();
        Dialog.addMessage("Sugg. : Sigma = 2");
        Dialog.addSlider("Sigma 2", 2, 30, 30);
        Dialog.addToSameRow();
        Dialog.addMessage("Sugg. : Sigma = 30");

        Dialog.show();


        //Saving options
        if(batch){
                inDir = Dialog.getString();
        }
        outDir = Dialog.getString();
        suffix = Dialog.getString();

        //Sigma parameters
        sigma1 = Dialog.getNumber();
        sigma2 = Dialog.getNumber();
}

//-------------------------------------------------------------------------
```

```
//Check if the image is in the right format, otherwise transforms it to 8-
bit
function checkFormat(){
        imgType = bitDepth();
        if (imgType == 24){
                run("8-bit");
        }
}

//---------------------------------------------------------------------------


//Duplicate the original images to keep it unprocessed, and generate images
for the difference of gaussians that need 2 images with different gaussian
blur sigma values.
function preProcessing(){
        ori = getTitle();
        run("Duplicate...", "title=img duplicate");
        img = getTitle();
        run("Duplicate...", "title=1 duplicate");
        img1 = getTitle();
        selectWindow(ori);
        run("Duplicate...", "title=2 duplicate");
        img2 = getTitle();
        selectWindow(ori);
        close();
}

//---------------------------------------------------------------------------


//Run the difference of gaussians on the previously duplicated images
function differenceOfGaussians(){
        selectWindow(img1);
        run("Gaussian Blur...", "sigma ="+sigma1);
        selectWindow(img2);
        run("Median...", "radius=2");
        run("Gaussian Blur...", "sigma="+sigma2);
        imageCalculator("Subtract create", "1","2");
        selectWindow("Result of 1");
        rename("Difference of Gaussians");
        DoG = getTitle();

        for (i = 1; i < 3; i++) {
                selectWindow(i); //Images for the difference of gaussians
are named 1 and 2, this "for" function closes both
                close();
        }
}

//---------------------------------------------------------------------------


//Run several modification on the image to run a skeletonize function, to
generate a properly segmented, binary image
function processingSegmentation(){
        run("Max...", "value=10000");
        run("Gamma...", "value=0.75");
        setAutoThreshold("Huang dark no-reset");
        run("Make Binary");
```

```
        rename("Binary Image");
        binary = getTitle();
        run("Skeletonize");
        run("Maximum...", "radius=1");
}

//---------------------------------------------------------------------------


//Allows the user to manually adjust the segmentation by drawing potential
missing cell walls, and generate the image for measurements
function manualSegmentation(){
        //Set up tools to manually adjust the segmentation process
        setForegroundColor(255, 255, 255);
        setBackgroundColor(0, 0, 0);
        waitForUser("Adjust the segmentation on Binary Image with brush
Tool\nToolset >> Drawing Tools");

        selectWindow(binary);
        resetThreshold();
        roiManager("Set Color", "black");
        roiManager("Set Line Width", 3);
        roiManager("Show All without labels");
        run("Flatten");
        run("8-bit");
        rename("Segmented");
        close(binary);
}

//---------------------------------------------------------------------------


//Draw a line along the epidermis and measure the plot profile
function measurement(){
        //Drawing a polyline for epidermis measurement
        setTool("polyline");
        setLineWidth(2);
        waitForUser("Draw a line along the epidermal
tissue");
        run("Plot Profile");
        run("Plots...", "list");
}

//---------------------------------------------------------------------------


//Save the plot profile list in txt format, which allows to open in excel
and measure the cells.
function saveData(){
        //Saving data
        oriSave = substring(ori, 0, lastIndexOf(ori, ".")); //Get the
original name of the image, without the extension
        selectWindow("Plot Values");
        save(outDir+"Plot Values_"+oriSave+".txt");
}

//---------------------------------------------------------------------------
```
